# Host phenology can select for multiple stable parasite virulence strategies

**DOI:** 10.1101/2021.06.08.447582

**Authors:** Hannelore MacDonald, Dustin Brisson

## Abstract

Host phenology is an important driver of parasite transmission dynamics and evolution. Prior research has demonstrated that host phenology can drive monocyclic, obligate-killer parasites to evolve an intermediate virulence strategy where all parasites kill their host just before the season ends to limit the death of parasite progeny in the environment. The impact of host seasonality on parasites that are not constrained to a monocyclic life-cycle, however, cannot be inferred from these results. Here we present a mathematical model that demonstrates that many, but not all, seasonal host activity patterns support multiple evolutionarily stable parasite strategies (ESS), although these strategies cannot coexist in the same system. The specific monocyclic and polycyclic parasite evolutionarily stable strategies in each phenological pattern are interspersed with less-fit mono- and polycyclic strategies (evolutionary repellors). The ESS that dominates each system at equilibrium is a function of the strategy of the parasite introduced into the system. The results demonstrate that host phenology can, in theory, maintain diverse parasite strategies among isolated geographic locations.

## Introduction

Classical ecological theory predicts that seasonality can increase niche space compared to equilibrial environments. In an equilibrial environment, slowly maturing species who efficiently collect resources (k-selected species) out-compete rapidly maturing species who are poor competitors (r-selected species).^1^ By contrast, seasonal resource fluctuations favor the high growth rate r-selected species when resources are temporarily abundant and the more competitive k-selected species when resources are temporarily scarce.^2–5^ The impact of seasonality on the vast diversity of parasite strategies observed in nature, however, remains equivocal.

Some studies have found evidence that seasonality generates diversity for certain parasites traits while other studies have been inconclusive. For example, an explicit trade-off between within-season transmission and between-season survival can select for both a polycyclic parasite strategy specialized on within-season infection initiation and a monocyclic parasite specialized on between-season survival.^6^ A similar model incorporating the same trade-off did not find evidence that seasonality can drive evolutionary branching,^7^ suggesting additional environmental conditions are important drivers. Seasonal host reproduction can also select for two divergent strategies including a latent strategy that can be reactivated later and an immediately transmissible strategy.^8^ While these studies show that seasonality can drive diversity in some specific situations, other studies have shown that seasonality does not impact the diversity of other important parasite traits such as virulence.^9^

Monocyclic and polycyclic parasites are subject to different life history constraints and selection pressures. For example, the time required to assemble parasite progeny constrains virulence evolution so that infected hosts are not killed before progeny are fully developed. Thus, parasites may be monocyclic in seasonal environments because there is only time to complete one infectious cycle. Parasites may also be constrained by mechanistic trade-offs where the number of progeny produced is mechanistically correlated with infection duration. Longer latency periods, the equivalent of lower virulence, can also be selectively advantageous in the absence of an explicit trade-off as low virulence monocyclic, obligate host-killer parasites remain in infected hosts to limit the decay of their progeny in the environment.^10^ However, parasites that are not constrained to a monocyclic lifestyle experience different selective pressures and may evolve alternate virulence traits. Prior investigations of obligate-killer parasites in non-seasonal environments suggest that high virulence traits are adaptive as they allow for exponential population growth.^11–14^ How host phenology impacts the strength and direction of selection pressures on polycyclic parasite strategies remains an open question.

Here we investigate the impact of host phenology on the virulence evolution of an obligate-killer parasite that is not constrained to the monocyclic life-style. We demonstrate that many host seasonality patterns can drive the evolution of both stable monocyclic and polycyclic strategies, although these strategies do not coexist. The evolutionary stable parasite strategy (ESS) in each system is a function of the virulence trait of the starting parasite population. Low virulence parasites evolve to a monocyclic ESS with a lower virulence trait, or a long latency period between infection and host death as seen previously,^10^ while high virulence parasites evolve to a polycyclic ESS with higher virulence to sequentially infect susceptible hosts within seasons. These results demonstrate that there are multiple evolutionary stable solutions for parasites in seasonal environments which provides clues for the evolutionary origins of monocyclic and polycyclic parasites.

### 1 Model description

The model describes the transmission dynamics of a free-living, obligate-killer parasite that infects a seasonally available host (Figure 1). *ŝ*(*n*) enter the system at the beginning of the season over a period given by the function *g*(*t, t_l_*). Hosts, *s*, have non-overlapping generations and are alive for one season. The parasite, *v*, must infect and kill the host to release new infectious progeny. The number of rounds of infection the parasite completes within a season depends on the parasite latency period length. If there is a long period between infection and progeny release, the parasite is monocyclic and completes one round of infection per season. If there is a short period between infection and progeny release, the parasite is polycyclic and can complete multiple rounds of infection per season.

**Figure 1:**
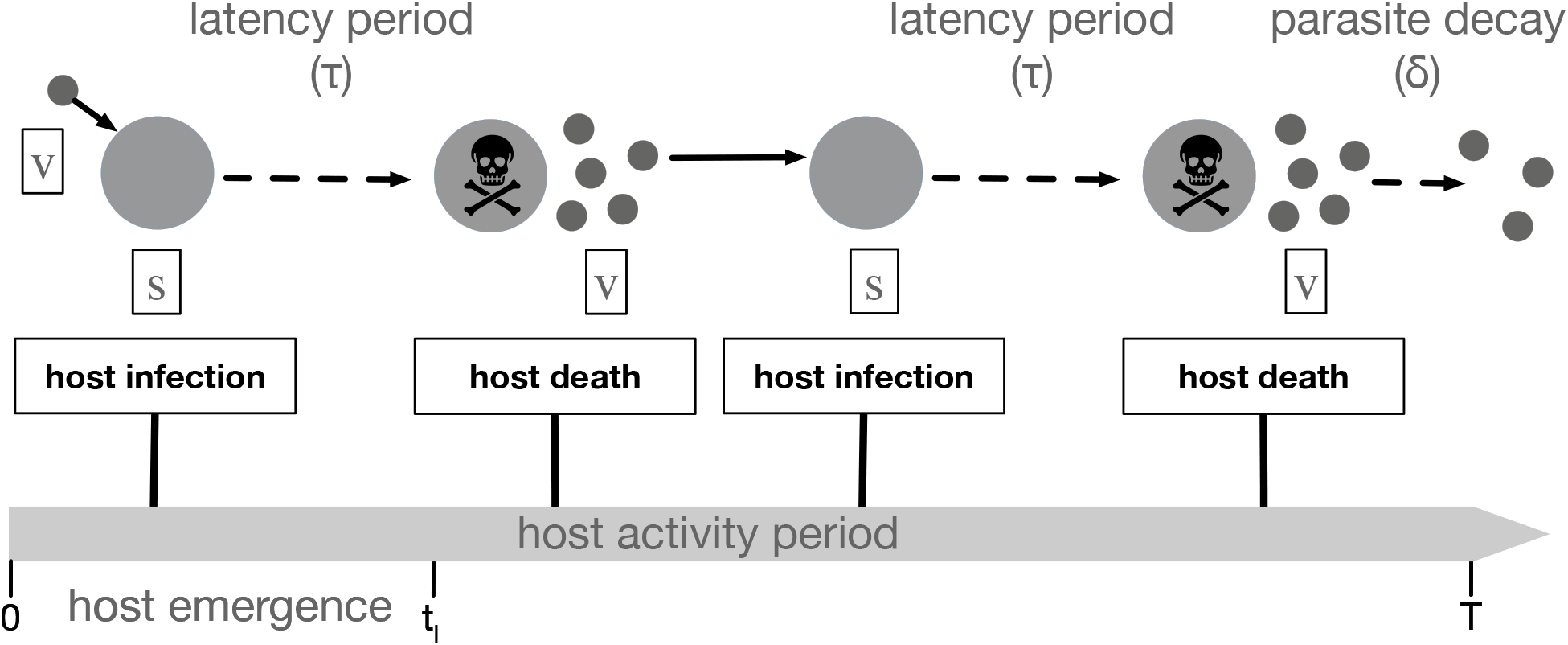
Diagrammatic representation of the infectious cycle within each season. All parasites (*v*) emerge at at the beginning of the season (*t* = 0) while all hosts (*s*) emerge at a constant rate between time *t* = 0 and *t* = *t_l_*. The rate of infection is density dependent such that the majority of the first round of infections occur near the beginning of the season when susceptible host and free parasite densities are high. Parasite-induced host death at time *τ* post-infection releases parasite progeny (*v*) into the environment where they decay in the environment from exposure at rate *δ*. If *τ* is short enough, more than one generation of infections can occur within the season. Parasite progeny that survive in the environment to the end of the season comprise the parasite population that emerge in the following season 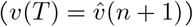.

The initial conditions in the beginning of the season are 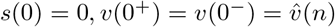 where 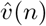 is the size of the starting parasite population introduced at the beginning of season *n* determined by the number of parasites produced in season *n* − 1. In some cases, the size of the emerging host cohort in season *n*, *ŝ*(*n*), is constant. We also explore the impact of host carryover between seasons by assuming that *ŝ*(*n*) is determined by the number of hosts that reproduced in season *n* − 1. The transmission dynamics in season *n* are given by the following system of delay differential equations (all parameters are described in Table 1):

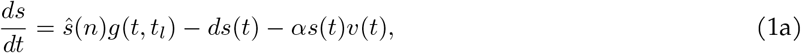

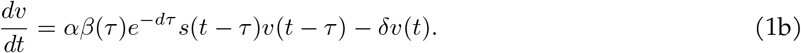

where *d* is the host death rate, *δ* is the decay rate of parasites in the environment, *α* is the transmission rate and *τ* is the delay between host infection and host death. *τ* is equivalent to virulence where low virulence parasites have long *τ* and high virulence parasites have short *τ*. *β* is the number of parasites produced upon host death. In most cases we assume *β* is a function of *τ*, *β*(*τ*) but also investigate the impact of a constant, trade-off free *β*.

**Table 1:** Model parameters and their respective values.

| Parameter | Description | Value |
| --- | --- | --- |
| $s$ | susceptible hosts | state variable |
| $v$ | parasites | state variable |
| $\hat{v}(n)$ | starting parasite population in season $n$ | state variable |
| $\hat{s}(n)$ | host cohort in season $n$ | state variable, $10^7$ when constant |
| $t_l$ | length of host emergence period | time (varies) |
| $T$ | season length | time (varies) |
| $\alpha$ | transmission rate | $10^{-8}/(\text{parasite} \times \text{time})$ |
| $\beta$ | number of parasites produced upon host death | 200 parasites when constant |
| $\delta$ | parasite decay rate in the environment | 2 parasites/parasite/time |
| $d$ | host death rate | 0.25 hosts/host/time |
| $\tau$ | latency period (1/virulence) | time (evolves) |
| $\sigma$ | host fecundity | 200 hosts |
| $\rho$ | density dependent parameter | 0.0001 |
| $b$ | trade-off parameter | 100 |

The function *g*(*t, t_l_*) is a probability density function that captures the per-capita host emergence rate by specifying the timing and length of host emergence. We use a uniform distribution (*U* (•)) for analytical tractability, but other distributions can be used.

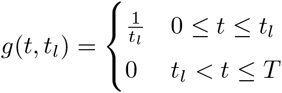

*t_l_* denotes the length of the host emergence period and *T* denotes the season length. The season begins (*t*_0_ = 0) with the emergence of the susceptible host cohort, *ŝ*. The host cohort emerges from 0 ≤ *t* ≤ *t_l_. ŝ*(*n*) is either constant or a function of the number of uninfected hosts remaining in the system at *t* = *T*. *v* parasites remaining in the system at *t* = *T* give rise to next season’s initial parasite population 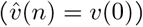.

Parasites that have not killed their host by the end of the season do not release progeny. Background mortality arises from predation or some other natural cause. We assume that infected hosts that die from background mortality do not release parasites because the parasites are either consumed or the latency period corresponds to the time necessary to develop viable progeny.^15, 16^

In previous work on a similar model we derived an analytical expression for parasite fitness as the density of parasites at the end of the season, *v*(*T*).^10^ However we cannot solve system (1) in the current framework analytically, thus all results were found by performing numerical computations.

#### 1.0.1 Between-season dynamics

To study the impact of the feedback between host demography and parasite fitness on parasite evolution we let the size of the emerging host cohort be a function of the number of uninfected hosts remaining at the end of the prior season

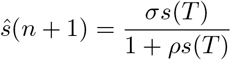

where *σ* is host reproduction and *ρ* is the density dependent parameter.

We have shown previously that host carryover generates a feedback between parasite fitness a nd host demography that can drive quasiperiodic dynamics for some parameter ranges.^17^

#### 1.0.2 Parasite evolution

To study how parasite traits adapt given different seasonal host activity patterns, we use evolutionary invasion analysis.^18, 19^ We first extend system (1) to follow the invasion dynamics a rare mutant parasite

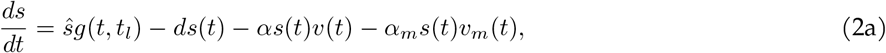

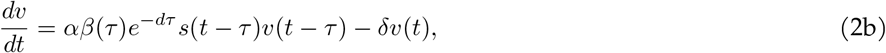

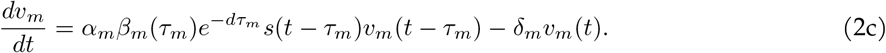

where *m* subscripts refer to the invading mutant parasite and its corresponding traits.

The invasion fitness of a rare mutant parasite depends on the density of *v_m_* produced by the end of the season (*v_m_*(*T*)) in the environment set by the resident parasite at equilibrium density 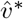. The mutant parasite invades in a given host phenological scenario if the density of *v_m_* produced by time *T* is greater than or equal to the initial *v_m_*(0) = 1 introduced at the start of the season (*v_m_*(*T*) ≥ 1).

To study the evolution of virulence traits (*τ*), we first assume all other resident and mutant traits are identical (*e.g. α* = *α_m_*). When *β* is a function of *τ*, we use assume that the number of new progeny released increases as the latency period increases: *β*(*τ*) = *b*(*τ* + 0.5)^0.8^. Note that when there is no trade-off between *β* and *τ*, the parasite growth rate in the host is essentially the trait under selection. That is, *β* is constant regardless of *τ*, thus the trait that is effectively evolving is the rate that new parasites are assembled in between infection and host death (*e.g.* long *τ* corresponds to slow assembly of new parasites.)

Virulence traits (*τ*) determine whether parasites are monocyclic or polycyclic by setting the latency period length. Highly virulent parasites have short latency periods that facilitate the completion of multiple rounds of infection per season. By contrast, low virulence parasites have long latency periods that only have time to complete one round of infection per season.

In previous work on similar models that only considered monocyclic parasites we were able to derive an analytical expression for mutant invasion fitness.^10, 1 7^ We are unable to solve the current model with polycyclic parasites analytically and instead determine parasite evolutionary endpoints numerically. As in previous analyses,^10, 17^ the invasion fitness of a rare mutant parasite depends on the density of *v_m_* produced by the end of the season (*v_m_*(*T*)) in the environment set by the resident parasite at equilibrium density 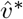. The mutant parasite invades in a given host phenological scenario if the density of *v_m_* produced by time *T* is greater than or equal to the initial *v_m_*(0) = 1 introduced at the start of the season (*v_m_*(*T*) ≥ 1). To find optimal virulence (*τ* *) for a given host phenological scenario, we find the uninvadable trait value numerically that satisfies

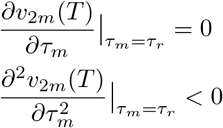

When hosts who survive to the end of the season (*s*(*T*)) reproduce to determine next season’s host cohort (*ŝ*), simulations are conducted to determine the outcome of parasite adaptation. Simulations are necessary because host carryover creates a feedback between parasite fitness and host demography that can drive cycling for some parameter ranges. When parasite-host dynamics are cycling, the density of *v_m_*(*T*) in the season the mutant was introduced does not reliably predict the outcome of parasite evolution as mutants with a selective advantage do not always invade.^17^ Simulations thus verify that the evolutionarily stable level of virulence is qualitatively the same as results when the emerging host cohort is constant across seasons and cycling cannot occur.

The simulation analysis was done by first numerically simulating system (1) with a monomorphic parasite population. A single mutant parasite is introduced at the beginning of the season after 100 seasons have passed. The mutant’s virulence strategy is drawn from a normal distribution whose mean is the value of *τ* from the resident strain. System (2) is then numerically simulated with the resident and mutant parasite. New mutants arise randomly after 1000 seasons have passed since the last mutant was introduced, at which point system (2) expands to follow the dynamics of the new parasites strain. This new mutant has a virulence strategy drawn from a normal distribution whose mean is the value of *τ* from whichever parasite strain has the highest density. System (2) continues to expand for each new mutant randomly introduced after at least 1000 seasons have passed. Any parasite whose density falls below 1 is considered extinct and is eliminated. Virulence evolves as the population of parasites with the adaptive strategy eventually invade and rise in density. Note that our simulations deviate from the adaptive dynamics literature in that new mutants can be introduced before earlier mutants have replaced the previous resident. Previous studies have shown that this approach is well suited to predicting evolutionary outcomes.^17, 20–22^

## Results

Monocyclic and polycyclic parasite strategies are both evolutionarily stable fitness optima in environments with seasonal host activity (Figure 2). However, monocyclic and polycyclic parasite subpopulations cannot coexist in the same host population. The evolutionary attractor that a population of parasites evolves towards is determined by their initial level of virulence, regardless of which strategy is the global optimum (Figure 2). Low fitness parasite strategies, which are evolutionary repellors, have virulence traits that kill hosts too quickly to limit progeny decay in the environment but do not kill hosts quickly enough to complete multiple infection cycles during the season.

**Figure 2:**
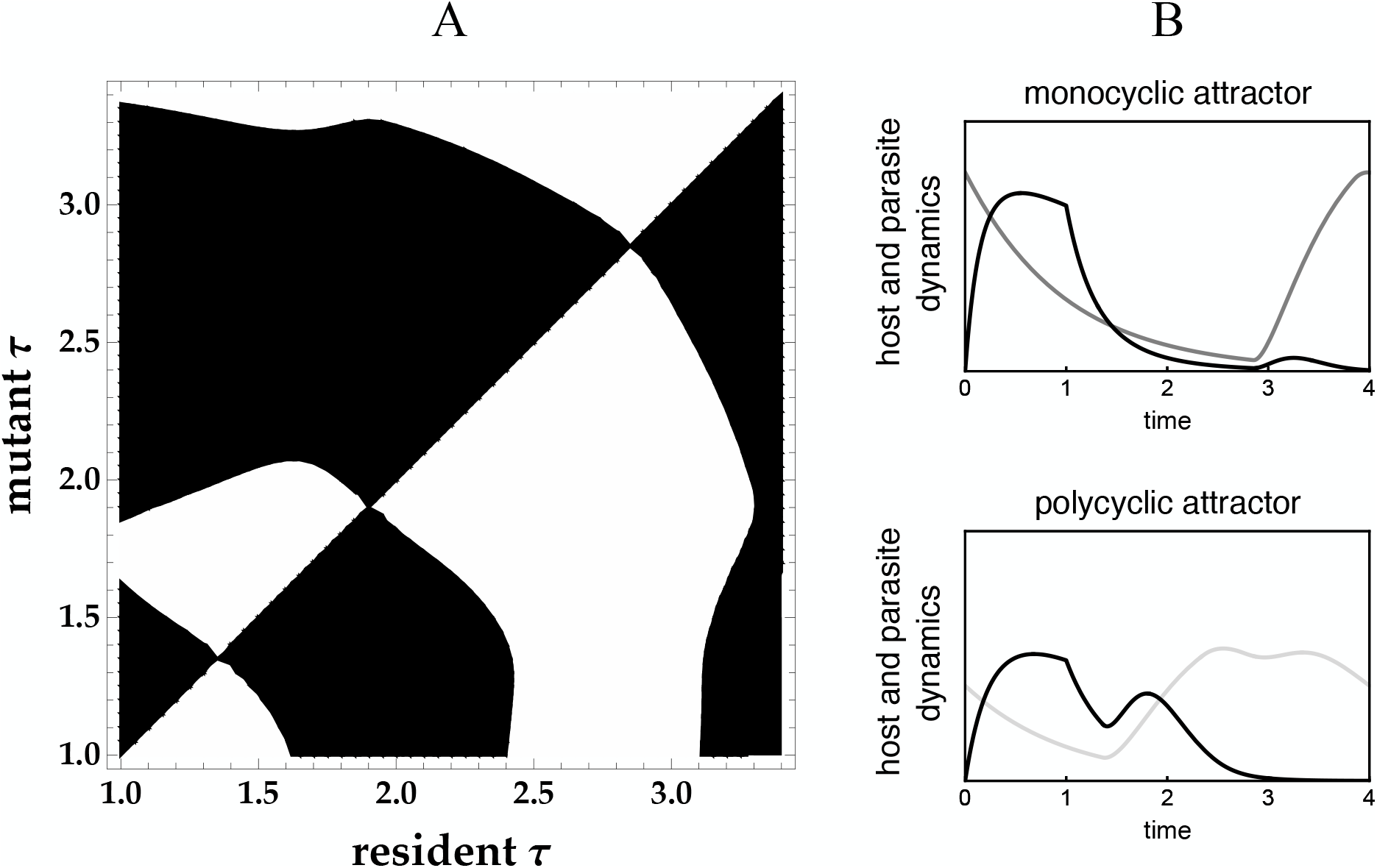
Seasonal host activity generates multiple parasite virulence attractors. **A.** Pairwise invasibility plot (PIP) shows the outcome of invasion by mutant parasite strains into resident parasite populations with virulence trait *τ*. Mutants possess an adaptive virulence trait and invade in black regions while they possess a maladaptive virulence trait and go extinct in white regions. The PIP shows two evolutionarily stable strategies (ESS) at *τ* ≈ 2.85 and *τ* ≈ 1.35 that are attractive and uninvasible. An evolutionary repellor lies between the two ESS at *τ* ≈ 1.9. **B.** The low virulence attractor (gray line, *τ* ≈ 2.85) releases new parasites just prior to the end of the season and is thus monocyclic. The high virulence attractor (light gray line, *τ* ≈ 1.35) is polycyclic and completes two generations of infections during the season for the parameter values shown here. Black line shows host dynamics. *T* = 4, *t_l_* = 1, *β*(*τ*) = *b*(*τ* + 0.5)^0.8^, all other parameters in Table 1. See Appendix A for an explanation of how the PIP was produced.

The progeny decay rate in the environment impacts the fitness of both monocyclic and polycyclic parasites. The virulence trait (latency period, *τ*) that optimizes fitness for both parasite strategies times the release of progeny from the final cycle of infected hosts to occur just prior to the end of the season. Parasites employing the monocyclic strategy evolve long latency periods to reach high densities at the end of each season by losing few progeny to environmental decay, as has been shown previously.^10^ The monocyclic strategy is the global optimum, resulting in the greatest number of progeny that survive to the end of the season, when (1) host death rates (*d*) are low such that infected hosts are unlikely to die prior to releasing progeny; (2) the variance in the timing that hosts become active during the season (*t_l_*) is small which results in a similarly small variance in the timing of infections near the start of the season; and (3) greater transmission rates (*α*) which similarly reduces the timing of infection variance.

The polycyclic strategies use higher virulence to complete multiple rounds of infection within a season. The polycyclic strategy is the global optimum, resulting in the greatest number of progeny surviving at the end of the season, when (1) host death rates (*d*) are high; and (2) the host emergence period (*t_l_*) is longer and the transmission rates (*α*) lower, both of which result in greater numbers of susceptible hosts later in the season who can be infected by progeny generated within the same season. Polycyclic parasites are particularly susceptible to self-shading, a process where the first cycle of infections reduce the number of susceptible hosts to the point where many of the first-cycle progeny fail to find a host within the season and are subject to decay in the environment.

The end of season density is not equivalent to invasion fitness for polycyclic parasites due to self shading. That is, high virulence polycyclic parasites invade and replace endemic lower virulence polycyclic parasites that achieve higher end of season densities. The higher virulence polycyclic parasites quickly kill infected hosts and progeny that infect many of the remaining susceptible hosts before the lower virulence parasite progeny have completed their first cycle. The progeny of the lower virulence parasites fail to find a susceptible host and decay in the environment even when their virulence trait would result in greater equilibrium densities in the absence of competition. Including a mechanistic trade-off between transmission and latency period length reduces the advantage of high virulence traits, who produce few progeny per infection with this trade-off, such that a moderately virulent polycyclic strategy is an ESS (Figure 2).

Shorter seasons and longer host emergence periods drive both the polycyclic and monocyclic optima toward higher virulence, similar to the results observed previously.^10^ More virulent monocyclic parasites successfully kill infected hosts and release progeny prior to the end of short seasons while more virulent polycyclic parasites can complete multiple infection cycles (Figure 3). Short host emergence periods result in simultaneously high host and parasite densities - and thus high density dependent infection rates - near the start of the season which favors less virulent parasites that kill hosts closer to the end of the season (Figure 3). However, the impact of host emergence periods on virulence evolution differs for the monocyclic and polycyclic optima. Small increases in the emergence period result in large virulence increases for the monocyclic parasite, but only minor increases in polycyclic parasite virulence, when the host emergence period length is short. This trend is reversed for long host emergence periods: small emergence period length increases cause negligible increases in monocyclic parasite virulence and large increases in polycyclic parasite virulence.

**Figure 3:**
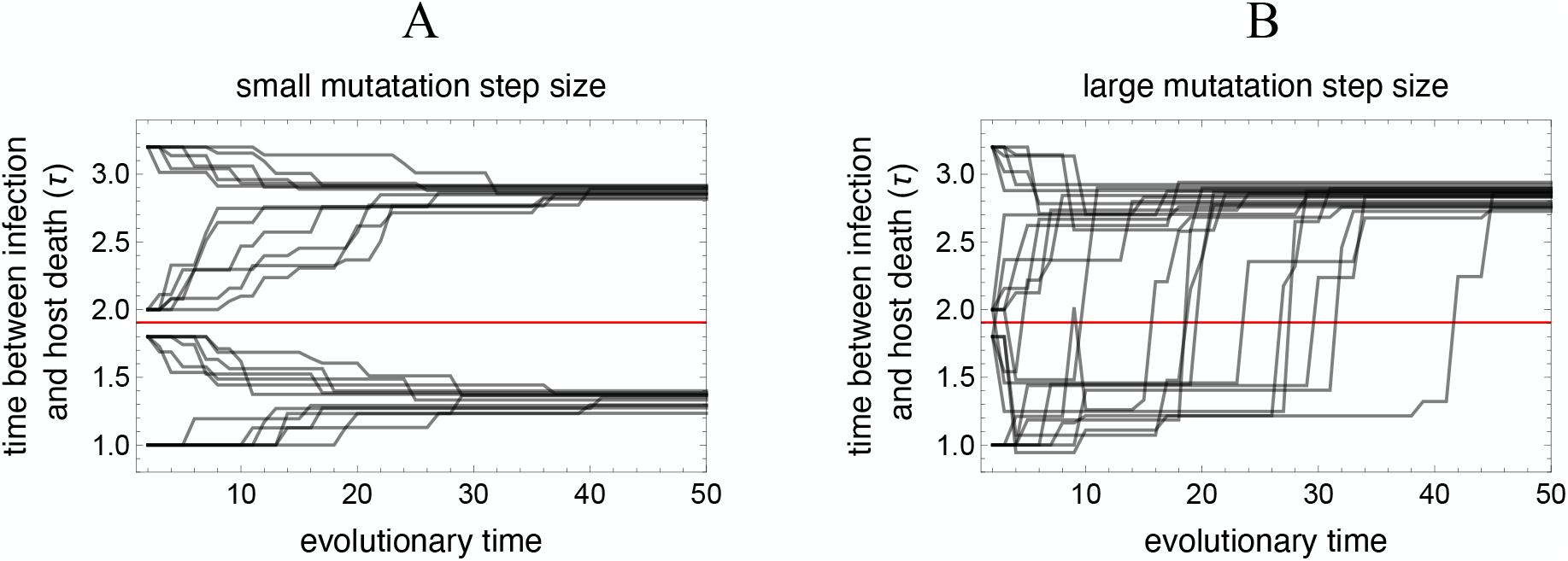
Initial conditions determine which virulence attractor parasite populations will evolve towards. A repellor exists between the two attractors at moderate virulence around *τ* = 1.9. (**A.**) If mutation step sizes are small, parasite populations with *τ* > 1.9 evolve towards the low virulence, monocyclic attractor at *τ* ≈ 2.85 while parasite populations with *τ* < 1.9 evolve towards the high virulence, polycyclic attractor at *τ*≈ 1.35. (**B.**) If mutation step sizes are large, all parasite populations eventually reach the low virulence, monocyclic attractor as this is the global optimum for these parameters. Plots show 24 independent simulation analyses with high or low mutation step sizes. Six runs start at *τ* = 3.2, *τ* = 2, *τ* = 1.8 and *τ* = 1, respectively. Evolutionary time represents the number of mutants introduced into each system. In a random season after at least 1000 seasons have passed since the last mutant was introduced, the parasite population with the highest density is set as the “resident” population and a new mutant is introduced with a virulence phenotype drawn from a normal distribution whose mean is the virulence phenotype of the “resident” parasite population. When the mutation step size is small: *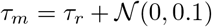*. When the mutation step size is large: *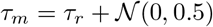*. Parameter values in this figure are identical to those in Figure 1.

Some environmental parameter states support only one parasite ESS (Figure 4). For example, monocyclic parasite fitness is limited in environments with high host mortality (*d*) as many infected individuals die prior to releasing parasites. Natural host mortality has little impact on the higher virulence polycyclic evolutionarily stable strategies that rapidly kill their hosts and infect subsequent hosts within a season. By contrast, the higher monocyclic parasite densities that result from lower host mortality increase early season incidence and leave few susceptible hosts for a second generation of polycyclic parasite infections. Thus, the low virulence, monocyclic ESS is the only viable evolutionary endpoint when host mortality is low.

**Figure 4:**
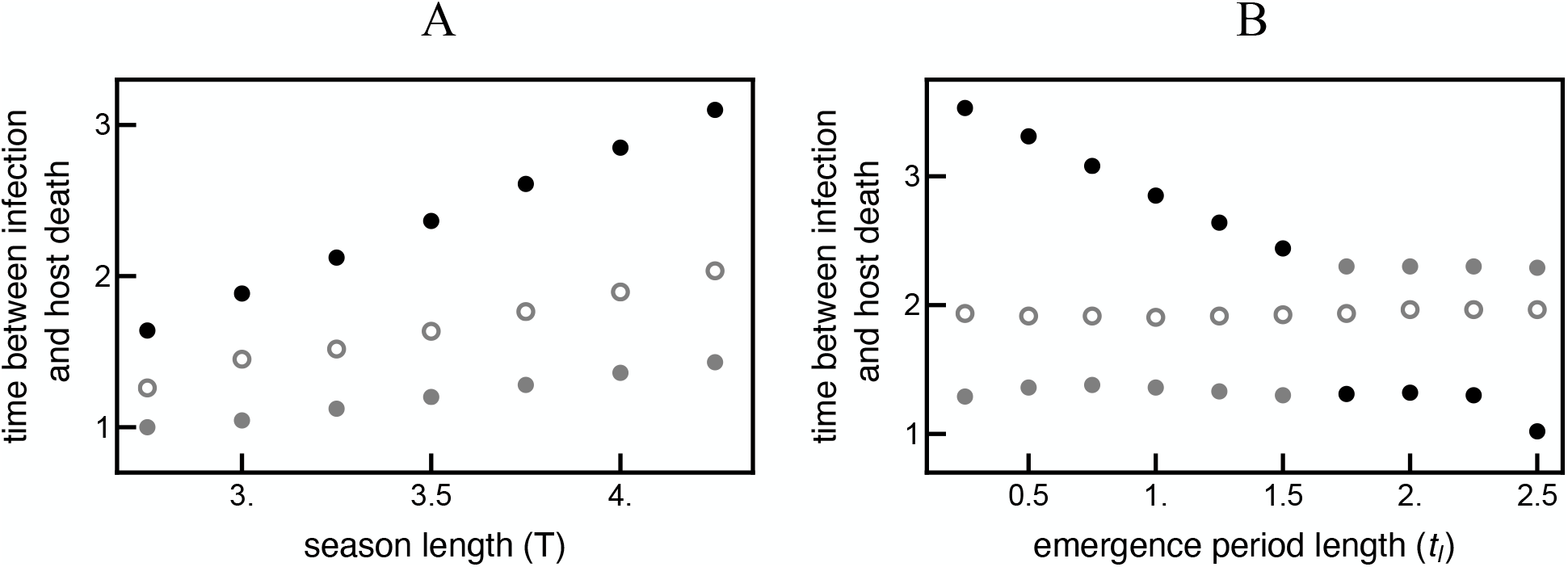
Host phenology impacts parasite virulence optimums. **A.** Longer seasons select for lower virulence for both the monocyclic and polycyclic attractors. **B.** Higher emergence variability selects for higher virulence for both the monocyclic and polycyclic attractors, however the impact of changing emergence period length is nonlinear. The strength *t_l_* has on the respective attractors varies: increases in *t_l_* when *t_l_* < 1.75 results in a large increase in virulence for the low virulence attractor but a small increase in virulence for the high virulence attractor while the opposite is true for *t_l_* > 1.75. Black points indicate global attractors, gray points indicate local attractors and hollow points indicate repellors. All other parameters are identical to those in Figure 1. See Appendix A for an explanation of how optimum virulence was found.

A feedback between host demography and parasite fitness can generate periodic host-parasite dynamics (Figure 5). This qualitative change in dynamical behavior does not impact the direction of virulence evolution. That is, the virulence level of the parasite originally introduced into the system will determine whether a polycyclic or monocyclic strategy evolves. However, the evolutionary rate proceeds much more slowly in dynamically cycling environments. Further, cyclic host-parasite dynamics routinely drive the polycyclic parasite population to extremely low densities while the monocyclic parasite population maintains densities well above extinction levels.

**Figure 5:**
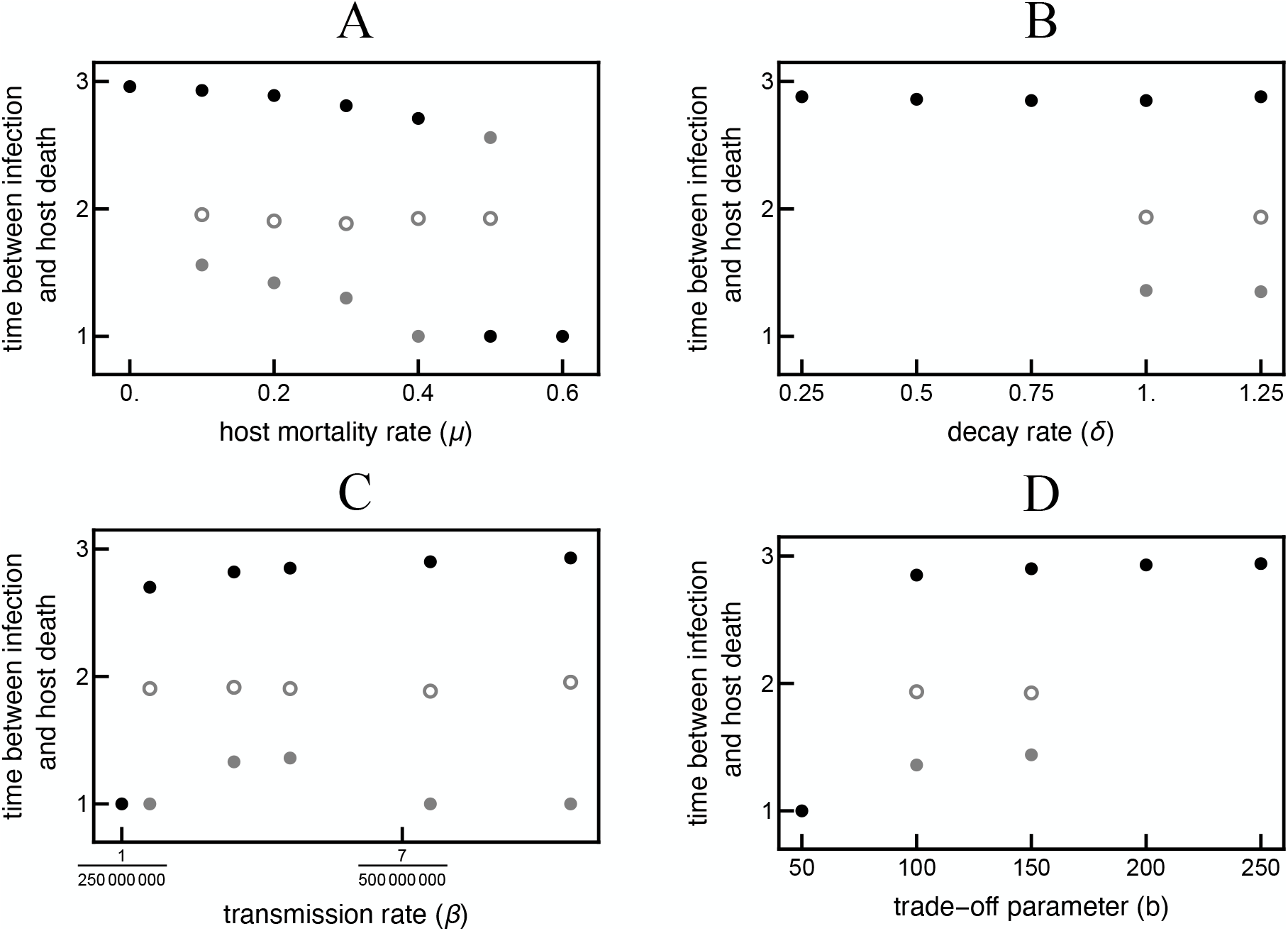
Certain conditions can destroy bistability or switch the global optimum. Parasite virulence attractors and repellors for changing: **A.** Host death rate, *μ* **B.** parasite decay rate (*δ*) **C.** transmission rate (*β*) **D.** trade-off parameter (*b*). **A.** Low natural host mortality (*μ*) drives high host densities and thus high early season incidence. High incidence early in the season selects for the low virulence, monocyclic strategy. High *μ* makes the high virulence, polycyclic attractor the global optimum as remaining in the host for long periods is risky. **B.** A low decay rate (*δ*) drives high parasite densities and thus high early season incidence. High incidence early in the season selects for the low virulence, monocyclic attractor. **C.** A low transmission rate (*β*) pushes the timing of infections to later in the season. Late infections select for the high virulence, polycyclic strategy as there is less time between infection and the end of the season. Higher *β* result in high early season incidence and thus drives both attractors towards lower virulence. **D.** Low values of the trade-off parameter (*b*) result in low parasite density and thus slow incidence. High virulence is adaptive when incidence is slow as parasites have less time to release progeny before the end of the season. High values of *b* result in high parasite density and thus high incidence early in the season. High early season incidence selects for the low virulence, monocyclic strategy. *T* = 4, *t_l_* = 1. When a parameter is not changing, its value is the same as in Table 1. See Appendix A for an explanation of how optimum virulence was found.

## Discussion

Many host phenological patterns can drive the evolution of multiple evolutionarily stable strategies. These phenological patterns support both a higher virulence strategy that completes multiple infection cycles within each season (polycyclic) and a lower virulence strategy that completes one infectious cycle each season (monocyclic). The monocyclic and polycyclic strategies are separated by an evolutionary repellor and are both evolutionarily stable attractors (Figure 4). Parasite populations that are more virulent than the repellor will evolve towards the polycyclic attractor while parasites that are less virulent than the repellor evolve towards the monocyclic attractor. Host seasonality is not predicted to permit the coexistence of diverse parasite strategies within the same host population but is predicted to drive diversity across geography.

The majority of prior studies suggest that obligate killer parasites maximize their fitness by rapidly killing and infecting new susceptible hosts, akin to the polycyclic strategy observed in the present model.^11–14^ This strategy relies on a constant supply of susceptible hosts which are temporally rare in seasonal environments. Parasites can limit the impact of seasonal absence of susceptible hosts by employing a less virulent, monocyclic strategy that coordinates the release of parasite progeny with the influx of susceptible hosts. In the current model, both strategies have a local fitness optimum in many phenological environments. Host seasonality is also predicted to maintain multiple evolutionarily stable strategies through mechanistic trade-offs between within-season transmission and between-season survival.^6^ In both cases, the monocyclic strategy evolves to decrease the impact of progeny decay in the environment during periods of host absence while the polycyclic strategy evolves to exponentially increase population sizes by rapidly exploiting hosts when they are abundant.

The parameter space that supports multiple evolutionarily stable parasite strategies in this model is broad but not universal. For example, the monocyclic strategy is the only evolutionary attractor in environments with low parasite decay rates. Low decay rates increase parasite densities which synchronizes early season incidence and decreases the number of susceptible hosts later in the season (self shading^23^) which limits reproductive success for polycyclic parasites. By contrast, high early season incidence increases monocyclic parasite fitness by synchronizing progeny release near the end of the season. This result deviates from the “Curse of the Pharoah” hypothesis which predicts that low parasite decay rates select for high virulence by decreasing the risk of leaving hosts early.^24–26^ The key difference between the current framework and previous work is that the “Curse of the Pharoah” hypothesis does not incorporate the impact of host demographic feedbacks on parasite fitness and thus predicts that the short-term benefit of an exponential increase in infections favors high virulence parasites.

Monocyclic and polycyclic parasites in some host phenological environments can achieve densities that are sufficient to destabilize host-parasite dynamics and instigate population cycles as observed previously with strictly monocyclic parasites.^17^ Population cycles occur when (*a*) parasites over-exploit host populations resulting in few susceptible hosts in subsequent years (*b*) leading to a parasite population crash (*c*) that allows host populations to recover. While the current model could not be solved analytically, a similar model suggested that cycles are driven by a Neimark-Sacker bifurcation.^17^ Cycling does not qualitatively alter the number of evolutionarily stable strategies supported by an environment nor the direction of parasite evolution within that system. However, populations cycles often drive polycyclic parasites toward extremely low densities that are at high risk for stochastic extinction (Figure 6). Polycyclic parasites exploit a greater number of susceptible hosts throughout the season than do monocyclic parasites, resulting in very small host populations and few infections in subsequent seasons. Thus, population cycles may drive polycyclic parasites extinct while maintaining monocyclic parasites in natural conditions.

**Figure 6:**
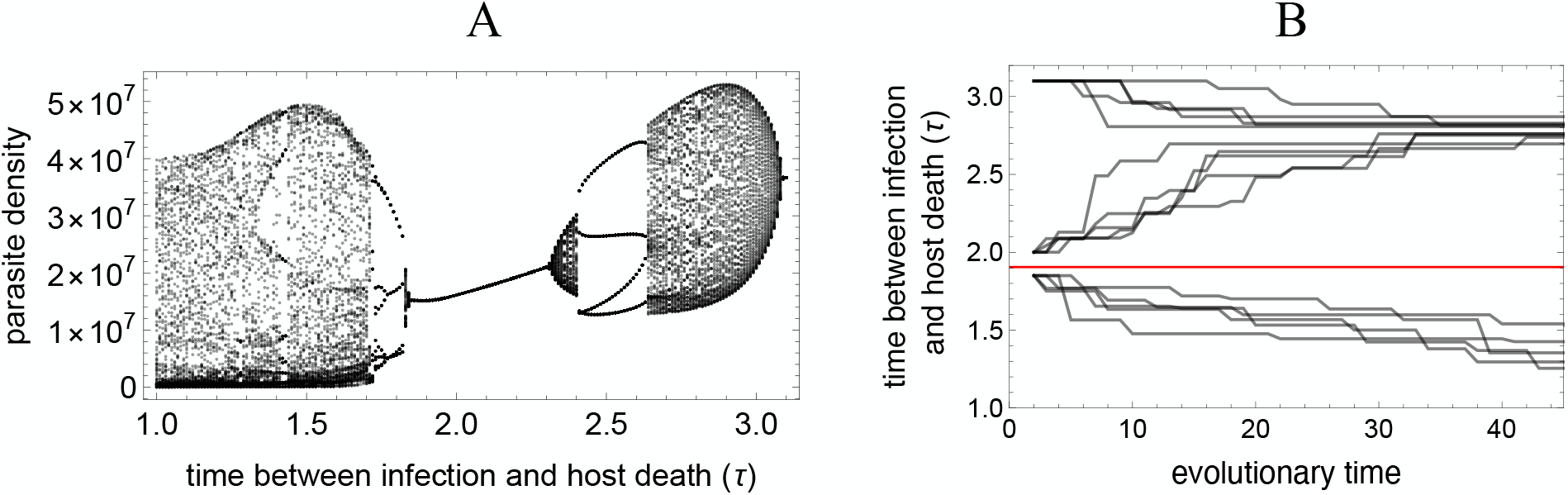
Virulence evolution generates periodic dynamics when host populations carryover from one season to the next. **A.** Parasite density increases as the virulence phenotype approaches the latency period (*τ*) that maximizes parasite fitness.^17^ Parasite populations can reach sufficiently high densities in some host phenological patterns to destabilize demographic dynamics resulting in a bifurcation that drives quasiperiodic parasite-host dynamics. The bifurcation diagram shows end of season parasite densities for parasites with different virulence phenotypes (*τ*) for seasons 800-900 in a system where the host season is short (*T* = 4) and hosts emerge synchronously (*t_l_* = 1). Moderate virulence parasites (1.85 < *τ* < 2.3) reach a stable equilibrium. The most fit polycyclic parasites with high virulence (*τ* < 1.85) and monocyclic parasites with low virulence (*τ* > 2.3) achieve densities that can disrupt dynamics and cause cycling. **B.** Periodic host-parasite dynamics do not qualitatively impact the evolutionary endpoints, *i.e.* high and low virulence attractors are separated by a repellor despite periodic dynamics. The same simulation analysis with small mutation step size was used as in Figure 2. *α* = 10^−7^, *b* = 75 to prevent polycyclic parasite extinction, all other parameters are the same as Table 1.

The primary conclusions of this model correspond with empirical data from some host-parasite study systems. For example, parasitic wasp species are more likely to employ a monocylic strategy at high latitudes where seasons are shorter and host emergence is more synchronous while the same species are dicyclic at low latitudes.^27–30^ Similarly, the monocyclic attractor is predicted to be the global optimum in short seasons and synchronous host emergence periods (Figure 4). However, experimental data are necessary to support host phenology as a driver of the evolution of monocyclic or polycyclic strategies across geography in this and other systems.

Several features of the current model can be altered to investigate more complex impacts of phenology on parasite diversity and virulence evolution. For example, relaxing the assumption that host populations reproduce once per season would likely favor higher virulence strategies as within-season host reproduction would reduce or eliminate the impact of self-shading for polycyclic parasites. The limited cost of self-shading in models with continuous host reproduction may explain why prior studies have not detected multiple evolutionarily stable parasite strategies for obligate killer parasites.

The strict obligate killer assumption can likely be relaxed without altering the result that many host phenological patterns support evolutionarily stable monocyclic or polycyclic strategies. For example, parasites that reduce host fecundity or increase host death rates upon progeny release likely experience similar evolutionary pressures on latency period duration as the obligate killer parasites modeled here. That is, releasing parasite progeny quickly (short latency period and high virulence) and releasing progeny near the end of the season (long latency period and low virulence) are both likely evolutionarily stable strategies. Many parasite-host systems conform to the assumptions of this model extension such as soil-borne plant pathogens, demicyclic rusts, post-harvest diseases, and many diseases systems infecting univoltine insects.^31–34^

Host phenology drives the timing and prevalence of transmission opportunities for parasites^35–42^ which impacts parasite life cycle strategies and virulence evolution.^8–10, 43, 44^ We add to this body of work by demonstrating that host phenology can also drive multiple evolutionarily stable parasite strategies. These results show that many host seasonal patterns impart selection pressures on parasites that can drive the evolution of parasite populations towards monocyclic and polycyclic life cycle strategies.

## Appendix A

In Appendix A we describe the numerical methods used to generate PIPs and to find evolutionary attractors and repellors. Code written for the numerical analysis is available upon request.

We follow the same approach as previous work^10, 17^ and define mutant invasion fitness as the density of mutant parasites produced by the end of the season (*v_m_*(*T*)) in the environment set by the resident parasite at equilibrium density 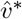. The mutant parasite invades if the density of *v_m_* produced by time *T* is greater than or equal to the initial *v_m_*(0) = 1 introduced at the start of the season (*v_m_*(*T*) ≥ 1).

It is not possible to derive an algebraic expression for mutant invasion fitness in this study as it was in previous studies. To generate PIP plots for pairs of resident’s with virulence (*τ_r_*) and mutant’s with virulence (*τ_m_*) we instead numerically find the density of *v_m_*(*T*) after one season in an environment set by *v_r_*. As in the previous analytical approach,^10, 17^ *v_m_*(*T*) = 1 corresponds to a neutral mutant, *v_m_*(*T*) > 1 corresponds to a mutant-resident pair in which the mutant parasite can invade and replace the resident, *v_m_*(*T*) < 1 corresponds to a mutant-resident pair that drives the mutant parasite extinct.

We use a similar approach to locate virulence trait values (*τ*) that correspond to evolutionary attractors and repellors. We again numerically find *v_m_*(*T*) after one season in an environment set by *v_r_*. Values of *τ* corresponding to attractors and repellors prevent small effect mutants with higher and lower virulence from invading. That is, when resident virulence is *τ_r_*, mutants with *τ_m_* = *τ_r_* + 0.01 and *τ_m_* = *τ_r_* − 0.01 cannot invade. We determine which points are attractors and repellors if there is more than one virulence trait value that prevents mutant invasion. Repellors are always found in between two attractors. To determine their location we find the value of *τ* in between the two attractors that corresponds to a minimum for *v_m_*(*T*).

To determine which attractor is the global attractor, we find the attractor that competitively excludes all others. Mutant parasites with the value of *τ* corresponding to the global attractor can invade a population of resident parasites with the value of *τ* corresponding to non-global, local attractors (*v_m_*(*T*) > 1). Resident parasites with the value of *τ* corresponding to the global attractor also prevent invasion of a mutant parasite with the value of *τ* corresponding to non-global, local attractors (*v_m_*(*T*) < 1).

## Notes

### Competing Interest Statement

The authors have declared no competing interest.

